# Mitochondrial Dysfunction and ER Stress in CB1 Receptor Antagonist-Induced Apoptosis in Human Neuroblastoma SH-SY5Y Cells

**DOI:** 10.1101/2024.10.21.619541

**Authors:** Kazuaki Mori, Akinobu Togo, Kota Yamashita, Taishi Umezawa, Keisuke Ohta, Toru Asahi, Chihiro Nozaki, Kosuke Kataoka

**Author notes:** Correspondence: Kosuke Kataoka, Graduate School of Engineering, Tokyo University of Agriculture and Technology, Tokyo, Japan. **Abbreviations** ATF4, activating transcription factor 4; CB1R, cannabinoid receptor type 1; CHOP, C/EBP homologous protein; eIF2α, eukaryotic translation initiation factor 2 α subunit; EIF2AK3, eukaryotic translation initiation factor 2-alpha kinase 3; ER, endoplasmic reticulum; ERO1α, ER oxidoreductin 1α; HAMMOC, hydroxy acid-modified metal oxide chromatography; Hp, Hemopressin; IP3R1, inositol-1,4,5-trisphosphate receptor 1; mPTP, mitochondrial permeability transition pore; mtCB1R, mitochondrial CB1R; PARP, poly ADP-ribose polymerase; PERK, protein kinase R (PKR)-like endoplasmic reticulum kinase; Rim, Rimonabant; SEM, scanning electron microscope; TRPV1, transient receptor potential channel vanilloid subfamily member 1.

## Abstract

Cannabinoid receptor type 1 (CB1R) is the key modulator of neuronal viability. In particular, CB1R antagonists provide neuroprotective effects on neurotoxicity caused by e.g. neuronal injury. However, the underlying mechanisms and potential limitations of CB1R antagonism remain unclear. Here we investigated the impact of environmental conditions on CB1R antagonist effects. Unexpectedly, we have found that cell-permeable CB1R antagonists, rimonabant and AM251, induced significant cell death in human neuroblastoma SH-SY5Y cells under serum-free conditions. Mitochondrial morphological analysis revealed mitochondrial swelling characterized by their network fragmentation and cristae reduction. Additionally, phosphoproteomics analysis showed an upregulation of phosphorylated EIF2AK3 (also known as PERK), leading to the activation of the eIF2α/ATF4/CHOP pathway, indicative of endoplasmic reticulum (ER) stress. Rimonabant and AM251 treatment also triggered caspase-dependent apoptosis, evidenced by the detection of cleaved caspase 3 and cleaved PARP. These results suggest that cell-permeable CB1R antagonists promote apoptosis via mitochondrial dysfunction and ER stress under serum-free conditions in SH-SY5Y cells. Our findings indicate that while CB1R antagonists may be neuroprotective in certain conditions, they may also pose a neurotoxic risk in environments characterized by cellular stress or nutrient deprivation.

## Introduction

Cannabinoid receptor type 1 (CB1R) is one of the key components of the endocannabinoid system, which maintains homeostasis in the body (Di Marzo et al., 2014). CB1R is primarily expressed in the central nervous system and is one of the most abundant G-protein-coupled receptors in the brain (Herkenham et al., 1990). CB1R signaling is crucial in neuronal viability and function (Kataoka et al., 2020; Zawilska et al., 2021). While the physiological activity of CB1R is generally associated with neuroprotection, excessive or prolonged activation can lead to adverse effects. For instance, acute exposure to delta-9 tetrahydrocannabinol (THC), a potent CB1R agonist, has been implicated in hippocampal neurotoxicity and cognitive deficits associated with cannabis use disorder (Chan et al., 1998; Lichtman et al., 2002). Furthermore, CB1R agonists have been shown to induce apoptosis in neuronal cells (Pasquariello et al., 2009; Tomiyama & Funada, 2014). The complex effects of CB1R signaling on neuronal health highlight the importance of maintaining a delicate balance in the endocannabinoid system.

CB1R antagonists have demonstrated neuroprotective effects in various contexts (Chan et al., 1998; Pegorini et al., 2006; Pasquariello et al., 2009; Liu et al., 2010; Tomiyama & Funada, 2014). These compounds have been found to prevent neurotoxicity caused by excessive CB1R activation, as evidenced by studies using rimonabant and AM251 (Chan et al., 1998; Pasquariello et al., 2009; Tomiyama & Funada, 2014). For instance, rimonabant inhibited THC-induced hippocampal neuron death and rescued ER stress-associated apoptosis in neuroblastoma cells (Chan et al., 1998; Pasquariello et al., 2009). Beyond counteracting CB1R agonist-induced toxicity, CB1R antagonists have shown promise in models of neurological injury, such as cerebral ischemia and diabetic neuropathy (Pegorini et al., 2006; Liu et al., 2010). These findings collectively suggest that CB1R antagonism may play a neuroprotective role by modulating CB1R signaling and potentially through other mechanisms.

Despite the growing evidence supporting the neuroprotective effects of CB1R antagonists, the underlying mechanisms and potential limitations of their action remain incompletely understood. The cellular environment plays a crucial role in determining the outcomes of pharmacological interventions (Zhang et al., 2016; Soutar et al., 2019; Gotten et al., 2022), yet the impact of environmental conditions on CB1R antagonist effects has not been fully explored.

In the present study, we have found unexpected neurotoxicity induced by CB1R antagonists rimonabant and AM251 under serum-free conditions in the SH-SY5Y cells but not in growth medium-supplemented conditions. Here, we investigated the mechanisms underlying this condition-dependent cell death induced by rimonabant and AM251. Our results demonstrate that these CB1R antagonists trigger mitochondrial damage and ER stress in SH-SY5Y cells under serum-free conditions, ultimately leading to caspase-dependent apoptosis. This finding challenges the conventional understanding of CB1R antagonism and highlights the importance of cellular context in determining drug effects.

## Materials and Methods

### Drugs

Rimonabant and AM251 were purchased from Selleck Chemicals (Texas, U.S.). Hemopressin was purchased from Tocris Bioscience (Minnesota, U.S.). Rimonabant and AM251 were dissolved in DMSO to 10 mM, and hemopressin was dissolved in Milli-Q water to 10 mM.

### Materials

Dulbecco’s modified Eagle’s medium (DMEM) with Low Glucose, penicillin-streptomycin (PS), and protease inhibitor cocktail set III, RIPA Buffer, PVDF membrane, polyoxyethylene(20) sorbitan monolaurate (Tween 20), and ImmunoStar LD were purchased from Fujifilm Wako (Osaka, Japan). Fetal bovine serum (FBS) was purchased from Cosmo Bio (Tokyo, Japan). LIVE/DEAD™ Viability/Cytotoxicity Kit was purchased from Thermo Fisher Scientific (Massachusetts, USA). VECTASHIELD Antifade Mounting Medium with DAPI was purchased from Vector Laboratories (California, USA). 16%-Paraformaldehyde aqueous solution was purchased from Nacalai Tesque (Kyoto, Japan). µ-Dish 35 mm Grid-500 (81166) was purchased from ibidi GmbH (Gräfelfing, Germany). 50% glutaraldehyde (3045) and epoxy resin (EPON812) were purchased from TAAB (Berkshire, England). Phosphatase inhibitor cocktail 2 and phosphatase inhibitor cocktail 3 were purchased from Sigma-Aldrich (St. Louis, USA). Sequencing grade modified Trypsin (Trypsin) was purchased from Promega (Wisconsin, USA). CDS Empore Disk C18 was purchased from CDS Analytical (Pennsylvania, USA). Can Get Signal® Immunoreaction Enhancer Solution was purchased from Toyobo (Osaka, Japan). Western BLoT Chemiluminescence HRP Substrate (Western BLoT) was purchased from Takara Bio (Shiga, Japan).

### Antibodies

Tom20 antibody (FL-145, sc-11415) was purchased from Santa Cruz Biotechnology (Texas, USA). The anti-rabbit fluorescent secondary antibody Alexa Fluor™ 488 (A-11008) was purchased from Thermo Fisher Scientific (Massachusetts, USA). Eukaryotic translation initiation factor 2 α subunit (eIF2α) antibody (11170-1-AP), phospho-eiF2α (peIF2α; Ser51) antibody, Activating transcription factor 4 (ATF4) antibody (10835-1-AP), and C/EBP homologous protein (CHOP) antibody (15204-1-AP) were purchased from Proteintech (Illinois, USA). Caspase 3 antibody (9662), poly ADP-ribose polymerase (PARP) antibody (9542), anti-rabbit HRP-linked antibody (7074), and anti-mouse HRP-linked antibody (7076) were purchased from Cell Signaling Technology (Massachusetts, USA). β actin antibody (010-27841) was purchased from Fujifilm Wako (Osaka, Japan).

### Cell lines and culture

Human neuroblastoma SH-SY5Y cell (RRID: CVCL_0019) was obtained from KAC Saibou.jp (Kyoto, Japan). SH-SY5Y cells were cultured in a growth medium (DMEM, 10% FBS, 1% PS) at 37 °C and 5% CO_2_.

### Live/Dead assay

SH-SY5Y cells were seeded at the density of 1.0 × 10^4^ cells/well in 100 μl of growth medium on 96-well clear plate. Three wells were used per condition in an independent experiment. 24 hours after cell seeding, serum starvation was conducted overnight using a serum-free medium (DMEM, 1% PS). The cells were treated with drugs in a growth medium or a serum-free medium for 24 or 12 hours at 37 °C and 5% CO_2_, respectively. Subsequently, a live/dead assay was conducted using the LIVE/DEAD™ Viability/Cytotoxicity Kit according to the manufacturer’s instructions. Fluorescent images were obtained at one place/well using an Eclipse Ti2-U fluorescence microscope (Nikon, Tokyo, Japan). The number of live cells was automatically counted using ImageJ Software (NIH, Maryland, USA). The executed macro was created with 8-bit, Convert to Mask, Watershed, and Analyze Particles (particle size threshold: 200 μm^2^).

### Immunofluorescence staining of mitochondrial morphology

SH-SY5Y cells were seeded at 5.0 × 10^4^ cells/well in 500 μl of growth medium on cover glass. As stated above, the cells underwent serum starvation and were treated with drugs in a serum-free medium or a growth medium for one hour. Subsequently, the cells were quickly washed with PBS and then fixed, permeabilized, and blocked according to our previous study (Mori et al., 2023). The cells were incubated with primary antibody Tom20 FL-145 (1:500) dissolved in PBS containing 0.3% BSA overnight at 4 °C. The next day, the cells were washed three times and incubated with secondary antibody Alexa Fluor™ 488 (1:1000) dissolved in PBS containing 0.3% BSA for 20 min at room temperature in the dark. After washing three times, the stained cells were mounted with VECTASHIELD Antifade Mounting Medium containing DAPI. Stained cell images were obtained using an FV1200 laser scanning microscope (Olympus, Tokyo, Japan).

### Mitochondrial morphology analysis in immunofluorescence images

Mitochondrial morphology was analyzed using ImageJ Software (NIH, Maryland, USA) according to previous studies (Valente et al., 2017; Chaudhry et al, 2020). The first, pre-processing (Unsharp Mask, Enhance Local Contrast, Median) was conducted on fluorescence images. Next, the images binarized by Make Binary. Subsequently, post-processing (Despeckle; Remove Outliners) was performed on the binarized images. Analyze Particles analyzed circularity of mitochondrion on the images. Finally, Skeletonize and Analyze Skeleton (2D/3D) was conducted on the images to analyze the average mitochondrial branch length. These processes were automated by the macro.

### Preparation of cell culture specimen for scanning electron microscope

SH-SY5Y cells were seeded at 4.0 × 10^4^ cells with 400 μl of growth medium on a 35 mm polymer bottom dish. As stated above, the cells underwent serum starvation and were treated with drugs in a serum-free medium for one hour. Subsequently, the cells were quickly washed with PBS and then fixed with pre-warmed fixative buffer (0.1 M cacodylate buffer, 100 mM NaCl, 2 mM CaCl_2_, pH 7.4, 2% paraformaldehyde, and 1% glutaraldehyde) for 15 min at room temperature. The fixed specimens were stored at 4 °C until postfixation by the osmium tetroxide-tannic acid-osmium tetroxide (OTO) method (Hirashima et al., 2022). Briefly, the specimens were rinsed five times with chilled 1 mM cacodylate buffer, then postfixed with 1% osmium tetroxide and 1.5% potassium ferrocyanide in 1 mM cacodylate buffer for one hour on ice. After five times washes with distilled water, the specimens were immersed in 1% thiocarbohydrazide solution for one hour at room temperature. They were then five times washed with distilled water and reacted with 1% osmium tetroxide solution for 30 min on ice. Following three times washes with distilled water, the specimens were dehydrated through an ethanol gradient (50% for 5 min on ice, 70% for 5 min on ice, 90% for 10 min on ice, and twice in 100% for 10 min at room temperature). The specimens were infiltrated by epoxy resin and then polymerized at 65 °C for two days. Resin blocks were then trimmed to 0.25mm^2^, and serial sections were cut using an ultramicrotome (ARTOS 3D, Leica, Wetzlar, Germany) using a diamond knife (Diatome, Biel, Switzerland). Serial sections were mounted on a silicon wafer and dried on a heating plate (60 °C); serial sections were double stained with uranyl acetate and lead acetate.

### Mitochondrial morphology analysis in SEM images

Electron micrographs of the cultured cells were obtained from their ultrathin sections as backscattered electron images using a scanning electron microscope (SEM; JSM-IT800; JEOL, Tokyo, Japan). Mitochondrial morphological characteristics (circularity, cross-sectional area) were measured using ImageJ Software. According to the previous study, cristae morphology classification was conducted double-blindly (Sun et al., 2007).

### Array tomography of the mitochondrial network

Approximately 100 consecutive images were obtained from 50 nm-thick ultrathin serial sections using SEM. The 3D shape of mitochondria was reconstructed from the 3D stacked image semimanually using Avizo 2024.1 (Thermo Fisher Scientific, Massachusetts, USA).

The mitochondrial complexity index (MCI) was employed as a quantitative measure of morphological complexity according to the previous report (Vincent et al., 2019). MCI was calculated using the formula:

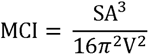

where SA is surface area and V is volume.

### Sample preparation for phosphoproteomic analyses

SH-SY5Y cells were seeded at 2.0 × 10^6^ cells/dish density in a growth medium on a 60 mm dish. As stated above, the cells underwent serum starvation were treated with drugs in a serum-free medium for one hour. Subsequently, protein extraction and digestion were performed according to our previous study with some modifications (Nakagami et al., 2019; Kamiyama et al., 2023). Briefly, total proteins were extracted in lysis buffer (10 mM Tris-HCl, pH 9.0, 8M urea, 2% phosphatase inhibitor cocktail 2, 2% phosphatase inhibitor cocktail 3, 2% protease inhibitor cocktail set III), followed by sonication and centrifugation at 12000 rpm for 10 min at 4°C. Crude extracts containing 320 μg of protein were subjected to the following enzymatic digestion. The solution was reduced with 10 mM dithiothreitol for 30 min and then alkylated with 50 mM iodoacetamide for 20 min in the dark. After 5-fold dilution with 50 mM ammonium bicarbonate in ultrapure water, proteins were digested with 8 μg trypsin per sample overnight at room temperature. Prepared digests were acidified with an equal volume of 2% trifluoroacetic acid to bind peptides to an in-house Stage tip made with Empore Disk C18. These acidified peptides were desalted with the Stage tip and dried using a centrifugal concentrator CC-105 (TOMY, Tokyo, Japan). Phosphopeptides were then enriched by the hydroxy acid-modified metal oxide chromatography (HAMMOC) method (Nakagami et al., 2019; Kamiyama et al., 2023), desalted again, and dried. Finally, the dried phosphopeptides were stored at -20°C until LC-MS/MS analysis.

### LC-MS/MS-based phosphoproteomics analyses

The dried phosphopeptides were dissolved in 20 μL of 2% acetonitrile containing 0.1% formic acid. According to the previous study (Kamiyama et al., 2023) , the dissolved peptides were directly loaded onto a C18 nano HPLC capillary column (NTCC-360/75-3, 75 µm ID×15 cm L, Tokyo, Nikkyo Technos) at 300 nL/min using an Easy-nLC 1200 (Thermo Scientific), and analyzed with Exploris 480 quadrupole-Orbitrap mass spectrometer (Thermo Fisher Scientific, Massachusetts, USA), equipped with an FAIMS Pro high-field asymmetric waveform aerodynamic ion mobility spectrometry device (Thermo Fisher Scientific, Massachusetts, USA). Peptide/Protein identification and MS1-based label-free quantification were carried out using Proteome Discoverer 2.5 (PD2.5, Thermo Fisher Scientific, Massachusetts, USA). MS2 spectra were searched with SEQUEST HT against the *Homoserines* protein database.

### Sample preparation for western blotting

SH-SY5Y cells were seeded at 2.0 × 10^6^ cells/dish density in a growth medium on a 60 mm dish. As stated, the cells underwent serum starvation were treated with drugs in a serum-free medium or a growth medium for three hours. After washing PBS, cells were immediately lysed using RIPA Buffer containing 1% protease inhibitor cocktail set III, 1% phosphatase inhibitor cocktail 2, and 1% phosphatase inhibitor cocktail 3. Subsequently, cell lysates were sonicated and then centrifuged at 16,000 g for 15 min at 4 °C. The supernatants were collected and mixed with an equal volume of 2X SDS sample buffer (125 mM Tris-HCl, pH 6.8, 4% sodium dodecyl sulfate (SDS), 10% sucrose, 0.01% Bromophenol Blue, 1% 2-mercaptoethanol). Samples were reduced for 5 min at 95 °C.

### Western blotting

10 μg of samples were loaded on SDS polyacrylamide gels. Following gel electrophoresis, the proteins were transferred to the PVDF membrane. Blocking was conducted using TBST (20 mM Tris-HCl, pH 7.6, 150 mM NaCl, 1% Tween 20) containing 5% skim milk at room temperature for one hour under shaking. Membranes were incubated with eIF2α (1:1000), peIF2α (1:1000), ATF4 (1:1000), CHOP (1:1000), caspase 3 antibody (1:1000), PARP antibody (1:1500), β actin antibody (1:10000) in Solution 1 of Can Get Signal at room temperature for one hour under shaking. The membranes were subsequently incubated with a secondary HRP antibody (1:5000) in Solution 2 of Can Get Signal at room temperature for one hour under shaking. Blots of membranes were detected as chemiluminescence by ImmunoStar LD for eIF2α, peIF2α, ATF4, CHOP, caspase 3, and PARP antibody or by Western BLoT for β actin antibody. Blot images were acquired using the ChemiDoc Touch MP Imaging System (Bio-Rad, California, USA). After blocking, primary antibody, and secondary antibody steps, membranes were washed with TBST at room temperature for 10 min 3 times. According to the previous report, membranes were incubated for sequential Western blotting of β actin antibody with 10% acetic acid to inactivate HRP chemiluminescence (Han et al., 2019).

### Data and statistical analysis

Kruskal-Wallis ANOVA with Dunn’s multiple comparisons and Mann-Whitney tests were used as statistical analysis. Statistical analysis was performed using GraphPad Prism 10.2.2–10.3.1 (California, USA).

## Results

### Cell death induced by cell-permeable CB1R antagonists under serum-free conditions in SH-SY5Y cells

To determine whether culture conditions alter the effects of CB1R antagonists on neuronal viability, human neuroblastoma SH-SY5Y cells were treated with varying doses of three types of CB1R antagonists in a growth medium or a serum-free medium (**Fig. 1a**): cell-permeable antagonists rimonabant and AM251, and a cell-impermeable antagonist hemopressin (Heimann et al., 2007). First, cell viability was assessed in cells treated with rimonabant, AM251, or hemopressin (100 nM–10 μM, respectively) in a growth medium. None of the antagonists affected cell viability in SH-SY5Y cells at any tested dose (**Fig. 1b**), suggesting that CB1R antagonists do not impact cell proliferation or induce neurotoxicity under normal growth conditions.

**Figure 1.**
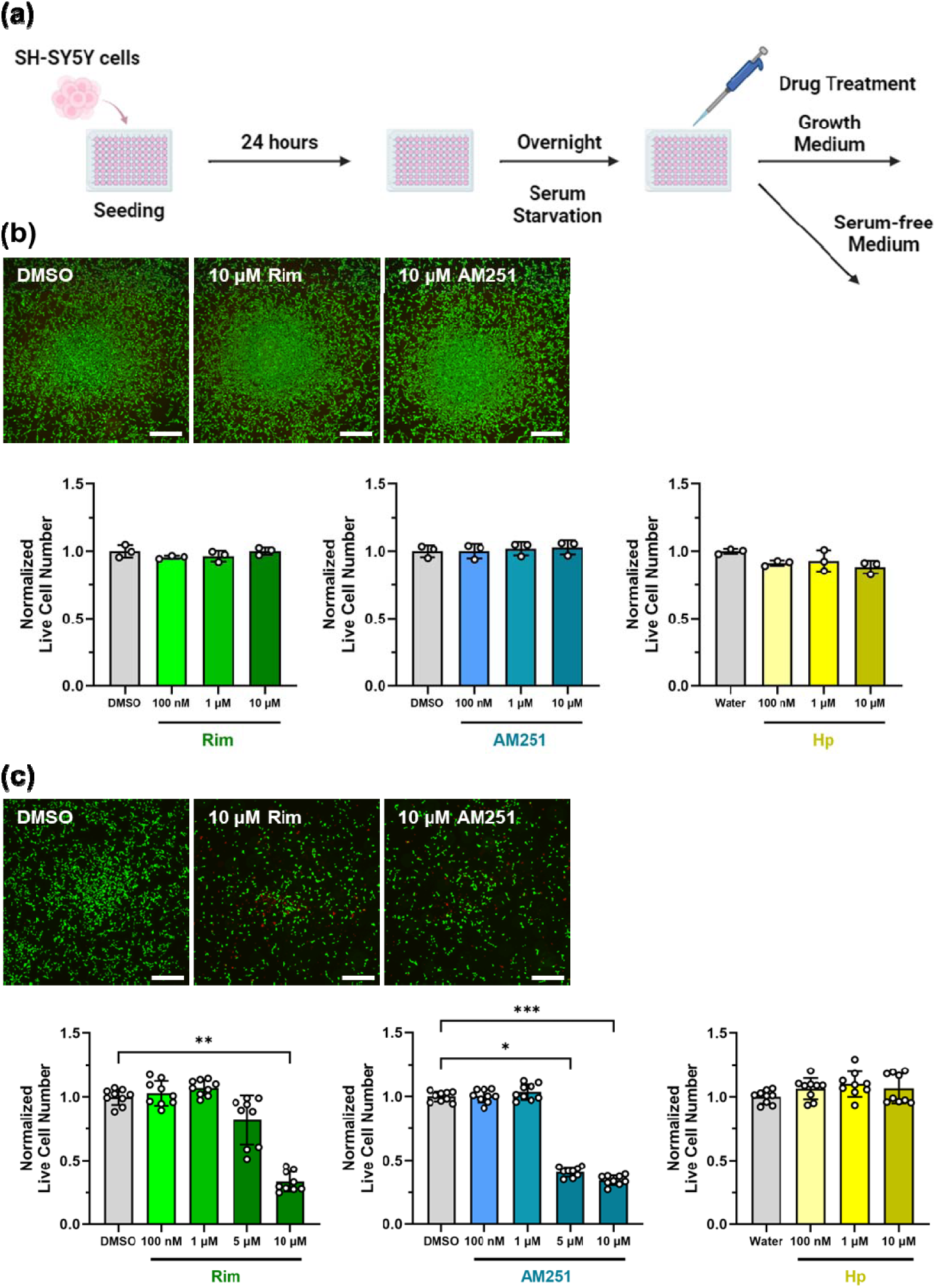
Live/Dead assay on SH-SY5Y cells treated with CB1R antagonists. (a) Experimental procedure of Live/Dead assay in s growth medium or a serum-free medium. (b) Cell viability in a growth medium. Cells were treated with rimonabant (Rim), AM251, or hemopressin (Hp) for 24 hours. Green, calcein-AM; red, ethidium homodimer-1. Scale bar: 500 μm. Calcein-AM positive cells are indicated as live cells and quantified by ImageJ software. The data of bar graph are shown as the mean ± SD for n = 3. Normalization was conducted based on the mean value of vehicle group. (c) Cell viability in a serum-free medium. Cells were treated with Rim, AM251, or Hp for 12 hours. The data of bar graph are shown as the mean ± SD for n = 9 (3 wells per independent experiment). Normalization was conducted based on the mean value of vehicle group per independent experiment. Data were analyzed by Kruskal-Wallis ANOVA with Dunn’s multiple comparison test. *, **, or *** indicate *P* < 0.05, 0.01, or 0.001, respectively.

Next, we examined the effects of these antagonists in a serum-free medium. Unexpectedly, 10 μM rimonabant decreased cell viability by 66% (**Fig. 1c**). AM251 also reduced viability by 59% at 5 μM and 66% at 10 μM (**Fig. 1c**). In contrast, hemopressin treatment did not induce cell death at any dose (**Fig. 1c**). These findings indicate that cell-permeable CB1R antagonists, but not cell-impermeable one, induce dose-dependent cell death under serum-free conditions in SH-SY5Y cells. Thus, the presence of serum in the medium is crucial in determining the neurotoxic potential of cell-permeable CB1R antagonists.

### Mitochondrial fragmentation induced by cell-permeable CB1R antagonists under serum-free conditions in SH-SY5Y cells

Mitochondria play a central role in cell death signaling (Green & Llambi, 2015). Their morphology changes in response to cellular stress, with fragmentation promoted under excessive stress to segregate damaged mitochondria from healthy ones (Youle & Van Der Bliek, 2012). To investigate the effect of CB1R antagonists on mitochondrial morphology under serum-free conditions, SH-SY5Y cells were treated with varying doses of rimonabant, AM251, or hemopressin in a serum-free medium for one hour, followed by immunofluorescence staining of mitochondria. At 10 μM, rimonabant significantly increased mitochondrial circularity and reduced branch length, indicating their fragmentation (**Fig. 2a**). Similarly, 10 μM AM251 increased comparable changes in mitochondrial morphology (**Fig. 2b**). In contrast, hemopressin had no effect on mitochondrial morphology at any tested dose (**Fig. 2c**). These findings suggest that cell-permeable CB1R antagonists cause dose-dependent mitochondrial fragmentation under serum-free conditions, while the cell-impermeable antagonist does not. This mitochondrial fragmentation may be a key factor contributing to the observed neurotoxicity of cell-permeable CB1R antagonists in serum-free environments.

**Figure 2.**
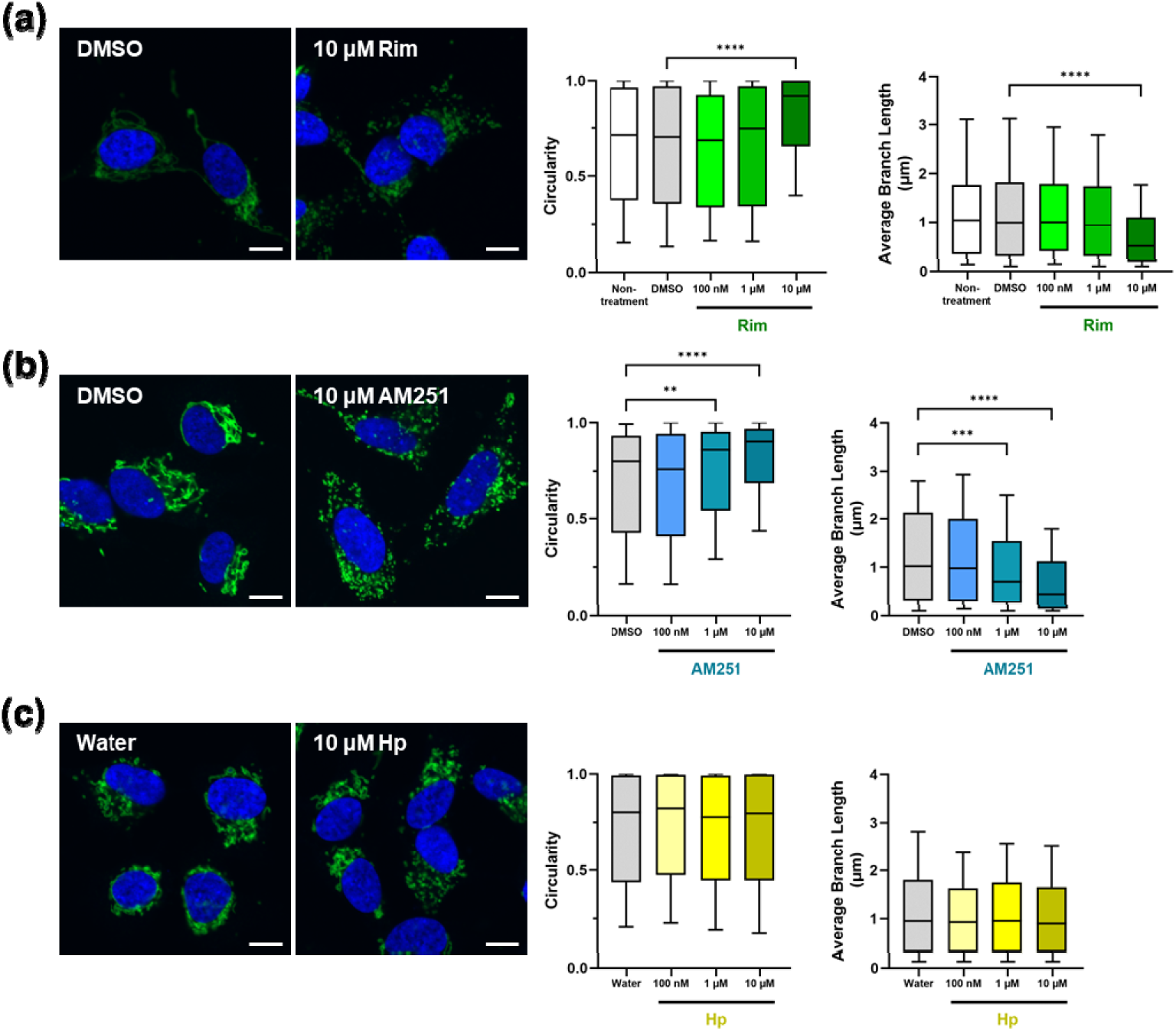
Mitochondrial morphology of SH-SY5Y cells treated with CB1R antagonists. Cells were treated with rimonabant (Rim, a), AM251 (b), or Hp (c) for one hour. Green, Tom20; blue, DAPI. Scale bar: 10 μm. Mitochondrial morphology was analyzed using ImageJ. Data were shown as Box and whiskers with 10–90 percentile. Rim: 51 cells (Non-treatment), 43 cells (DMSO), 51 cells (100 nM), 51 cells (1 μM), and 47 cells (10 μM); AM251: 46 cells (DMSO), 57 cells (100 nM), 49 cells (1 μM), and 35 cells (10 μM); Hp: 55 cells (Water), 56 cells (100 nM), 60 cells (1 μM), and 55 cells (10 μM). Data were analyzed by Kruskal-Wallis ANOVA with Dunn’s multiple comparison test. **, ***, or **** indicate *P* < 0.01, 0.001, or 0.0001 respectively.

We also examined the effects of cell-permeable CB1R antagonists on mitochondrial morphology under growth conditions. SH-SY5Y cells were treated with 10 μM rimonabant or 10 μM AM251 in a growth medium for one hour. Rimonabant showed no effect on mitochondrial morphology (**Fig. S1**), while AM251 slightly increased mitochondrial circularity and reduced branch length (**Fig. S1**). These results indicate that cell-permeable CB1R antagonists have only minor mitochondrial toxicity under growth conditions. In summary, high doses of these antagonists may exert mild neurotoxic effects, but the occurrence of cell death likely depends on environmental conditions.

### Mitochondrial swelling induced by cell-permeable CB1R antagonists under serum-free conditions in SH-SY5Y cells

To observe mitochondrial fragmentation induced by CB1R antagonists at higher resolution, we employed a scanning electron microscope (SEM). Cells were treated with 10 μM rimonabant, 10 μM AM251, or 10 μM hemopressin in a serum-free medium for one hour. As a result, both rimonabant and AM251 increased mitochondrion circularity and reduced mitochondrial area, whereas hemopressin had no effect on these parameters (**Fig. 3a, b**). These SEM observations corroborate the immunostaining results, confirming that cell-permeable CB1R antagonists induce mitochondrial fragmentation.

**Figure 3.**
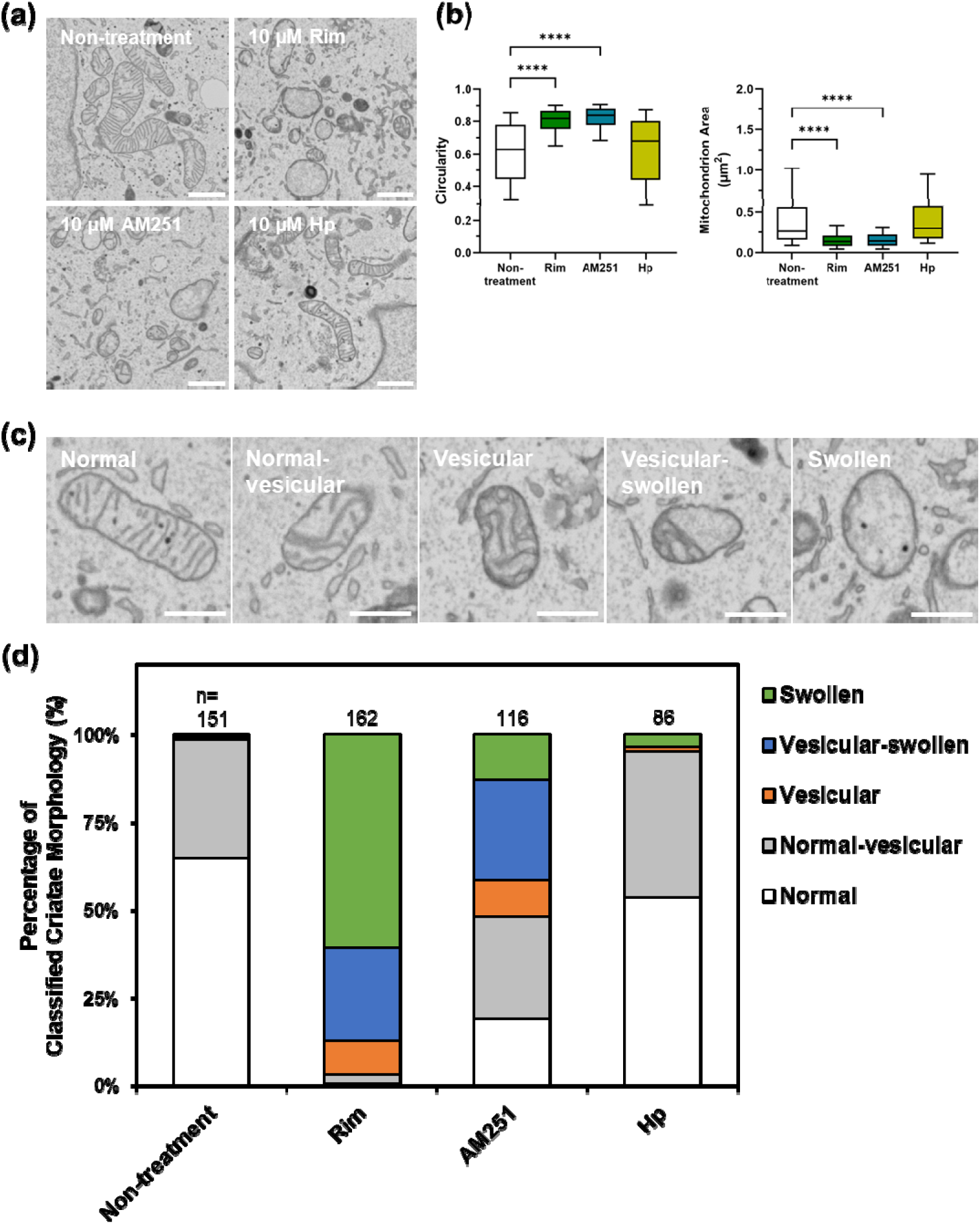
High-resolution observation of mitochondria morphology of SH-SY5Y cells treated with CB1R antagonists. (a) Scanning electron microscope images of mitochondria. Cells were treated with 10 μM of CB1R antagonists (rimonabant (Rim), AM251, or hemopressin (Hp)) for one hour, respectively. Scale bar: 1 μm. (b) Mitochondrial morphology analysis. Data were shown as Box and whiskers with 10–90 percentile. Nontreatment: 179 mitochondria; Rim: 348 mitochondria; AM251: 334 mitochondria; Hp: 187 mitochondria. Data were analyzed by Kruskal-Wallis ANOVA with Dunn’s multiple comparison test. **** indicates *P* < 0.0001. (c) Classification of cristae morphology by three stages: normal, normal–vesicular, vesicular, vesicular–swollen, and swollen. Scale bar: 500 nm. (d) Quantification of the overall cristae morphology. Cristae morphology was classified in a double-blind fashion. The number of total classified mitochondria were indicated at the top of the bar in each condition.

To analyze the effect of CB1R antagonists on mitochondrial cristae, we classified cristae morphology into five stages: normal, normal–vesicular, vesicular, vesicular–swollen, and swollen, according to the previous study (**Fig. 3c**) (Sun et al., 2007). In vesicular cristae, the inner membrane encloses a separate vesicular matrix, forming a continuous internal cavity. Swollen cristae are characterized by expanded matrix space with fewer cristae and ruptured outer membranes. In untreated conditions, 65% or 34% of cristae morphology was classified as normal or normal–vesicular, respectively (**Fig. 3d**). However, rimonabant treatment resulted in only 0.62% normal cristae with 66% swollen and 27% vesicular–swollen cristae (**Fig. 3d**). Similarly, AM251 treatment led to 13% swollen and 28% vesicular–swollen cristae (**Fig. 3d**). In contrast, hemopressin treatment maintained 53% normal and 41% normal–vesicular cristae morphology (**Fig. 3d**). These findings indicate that cell-permeable CB1R antagonists, but not cell-impermeable one, induce mitochondrial swelling under serum-free conditions in SH-SY5Y cells.

### 3D reconstruction of mitochondria treated with cell-permeable CB1R antagonists

To observe changes in the 3D mitochondrial network following treatment with cell-permeable CB1R antagonists, 3D reconstructions of mitochondria were conducted based on approximately 100 consecutive SEM images. All mitochondria in a single SH-SY5Y cell were reconstructed for both untreated and 10 μM rimonabant treated conditions (**Fig. 4a, SI Movie 1, 2**). Representative 3D mitochondrial morphologies were shown in zoomed images (**Fig. 4b**). 3D mitochondrial morphology was quantified by mitochondrial volume and mitochondrial complexity index (MCI) according to the previous report (Vincent et al., 2019). Rimonabant treatment significantly decreased both mitochondrial volume and MCI, indicating that mitochondria treated with rimonabant were fragmented and exhibited disrupted connectivity (**Fig. 4c**). Furthermore, a portion of the mitochondria treated with 10 μM AM251 was reconstructed (**Fig. 4d**). Similar to rimonabant treatment, the representative 3D image showed fragmented mitochondria with disrupted connectivity (**Fig. 4d**). Overall, these 3D reconstructions demonstrate that cell-permeable CB1R antagonists induced fragmentation of 3D mitochondrial network, corroborating our previous observations from 2D imaging.

**Figure 4.**
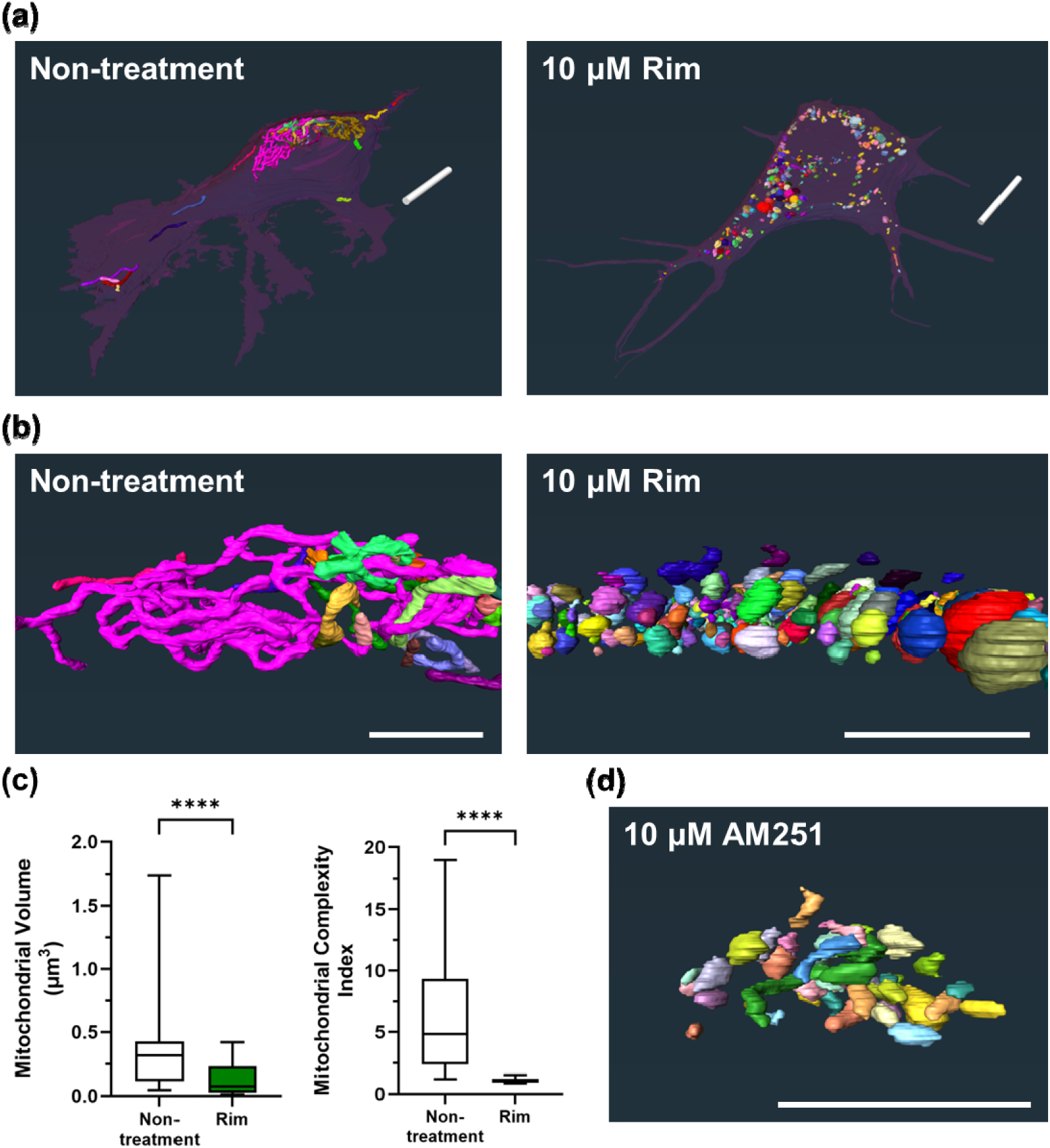
3D reconstruction of mitochondria using SEM array tomography. (a) Images of all mitochondria of single cell in non-treatment and 10 μM rimonabant (Rim) treatment conditions. Colored particles, mitochondria; reddish purple, cell. Scale bars: 10 μm. (b) Representative zoomed images of mitochondria in non-treatment and 10 μM Rim treatment conditions. Scale bars: 5 μm. (c) Mitochondrial network analyses. Data were shown as Box and whiskers with 10–90 percentile. Non-treatment: 35 mitochondria; Rim: 346 mitochondria. Data were analyzed by Mann-Whitney test. **** indicates *P* < 0.0001. (d) Representative image of mitochondria in 10 μM AM251 treatment condition. Scale bars: 5 μm.

### Molecular pathway related to cell death induced by cell-permeable CB1R antagonists under serum-free conditions in SH-SY5Y cells

To explore the molecular mechanisms underlying cell death induced by CB1R antagonists under serum-free conditions, we performed phosphoproteomics analyses. Phosphorylation is closely linked to protein activity and plays a critical role in regulating signaling pathways, especially those involved in programmed cell death (Lei & Davis, 2003). SH-SY5Y cells were treated with 10 μM rimonabant or 10 μM AM251 in a serum-free medium for one hour. Phosphopeptides were enriched using the hydroxy acid-modified metal oxide chromatography (HAMMOC) method (Nakagami et al., 2019; Kamiyama et al., 2023). As a result, phosphorylation of Ser715 in EIF2AK3 (eukaryotic translation initiation factor 2-alpha (eIF2α) kinase 3, also known as PERK) was significantly up-regulated in both rimonabant- and AM251-treated cells (**Fig. 5a, b**). EIF2AK3 is a main signal transducer of endoplasmic reticulum (ER) stress (Bertolotti et al., 2000).

**Figure 5.**
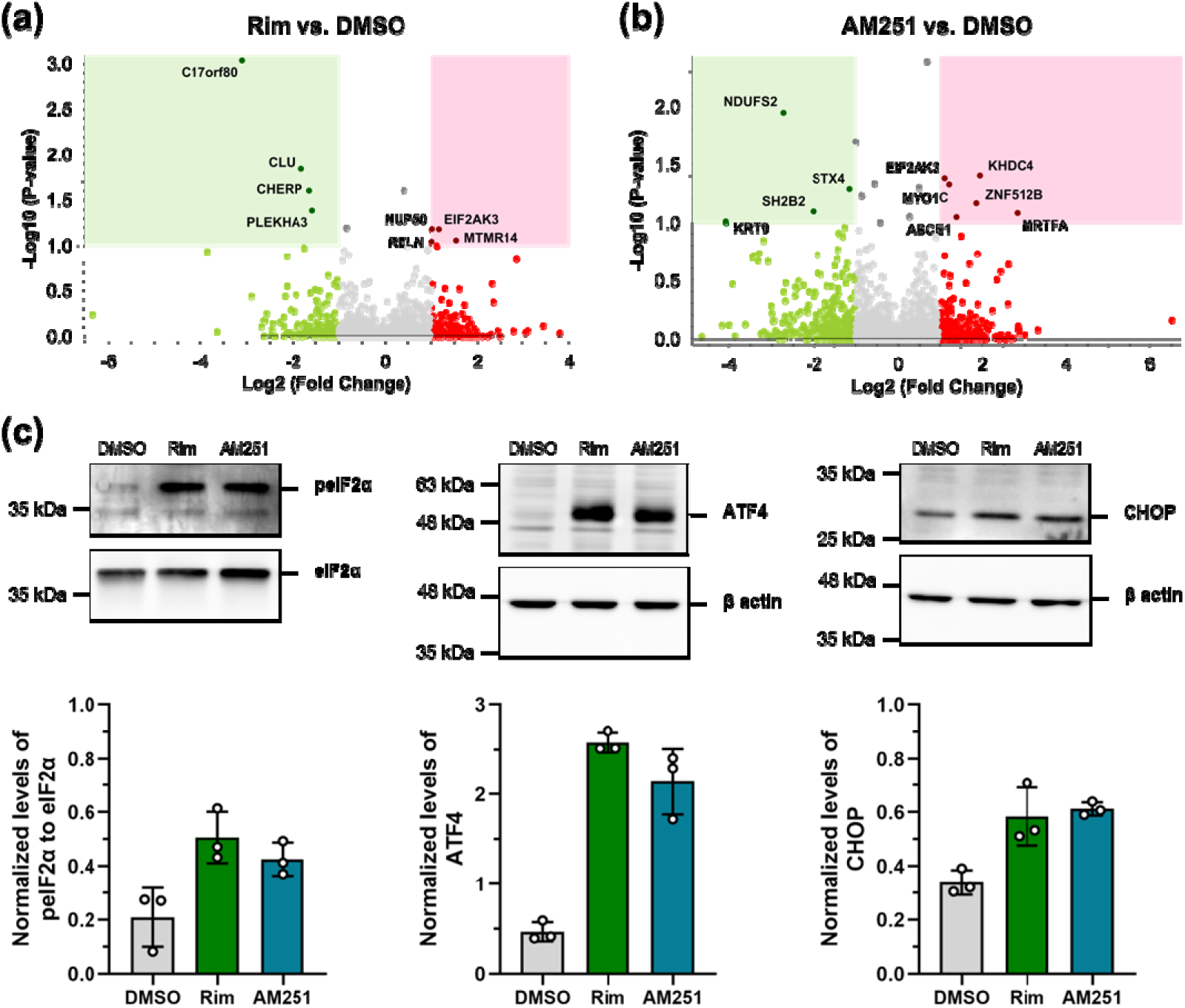
Investigation of molecular pathway of cell death induced by CB1R antagonists in SH-SY5Y cells. (a,b) Volcano plots showing phosphoproteomic analyses. Lysates of cells treated with 10 μM rimonabant (Rim, a) or 10 μM AM251 (b) for one hour were used. The averages of the phosphoproteomic expression data of Rim or AM251 treatment group (N = 3) were compared with the averages of the data for vehicle treatment group (N = 3). P-value was calculated by performing t test of PD 2.5. P-value setting is 0.1, and the Log 2 (Fold Change) setting is 1. Red circles show phosphosites which have significant increases. Green circles show phosphosites which have significant decreases. Gray circles are phosphosites without any differences. (c) Western blotting on EIF2AK3 downstream signaling pathway; phospho-eIF2α (peIF2α)/ATF4/CHOP. Lysates of cells treated with 10 μM Rim or 10 μM AM251 for three hours were used. The data of bar graph are shown as the mean ± SD for n = 3 (3 per independent experiment). Expression levels of peIF2α was normalized by eIF2α, and ATF4 and CHOP were normalized by β actin expression per independent experiment.

Phosphorylation of Ser715 PERK activates phosphorylation of Ser51 eIF2α (peIF2α) (Harder et al., 2013), triggering a pro-apoptotic signaling cascade involving activating transcription factor 4 (ATF4) and C/EBP homologous protein (CHOP) (Bertolotti et al., 2000; Gundu et al., 2022). To confirm the activation of this pathway, we examined the expression of peIF2α, ATF4, and CHOP in SH-SY5Y cells treated with 10 μM rimonabant or 10 μM AM251 in a serum-free medium for three hours. Both treatments resulted in increased expression of peIF2α, ATF4, and CHOP (**Fig. 5c**). In summary, these results indicate that cell-permeable CB1R antagonists activate the EIF2AK3/eIF2α/ATF4/CHOP axis, suggesting ER stress-mediated apoptosis.

We also examined the expression of peIF2α, ATF4, and CHOP in treatment of 10 μM rimonabant or 10 μM AM251 under growth conditions. SH-SY5Y cells were treated with 10 μM rimonabant or 10 μM AM251 in a growth medium for three hours. Neither treatment affected the expression of peIF2α, ATF4, and CHOP (**Fig. S2**). These findings suggest that under growth conditions, cell-permeable CB1R antagonists do not induce ER stress or exhibit toxicity toward ER.

### Apoptosis induced by CB1R antagonists under serum-free conditions in SH-SY5Y cells

To determine whether CB1R antagonists induce apoptosis under serum-free conditions, we analyzed the expression of apoptosis markers, including cleaved caspase-3 and cleaved poly ADP-ribose polymerase (PARP). Caspase 3 is a key executioner of apoptosis (Boatright & Salvesen, 2022), which, when cleaved, exposes its catalytic site and enables the cleavage of various substrates, including DNA fragmentation factors and cytoskeletal components (Wright et al., 2006; Boatright & Salvesen, 2022). PARP, a substrate of caspase 3, is cleaved into 89 kDa and 24 kDa fragments during apoptosis (Mashimo et al., 2021). SH-SY5Y cells were treated with 10 μM rimonabant or 10 μM AM251 in a serum-free medium for three hours. Western blot analysis revealed the presence of cleaved caspase 3 (**Fig. 6a**) and the 89 kDa cleaved PARP fragment (**Fig. 6b**) in both rimonabant- and AM251-treated cells. These findings indicate that cell-permeable CB1R antagonists induce caspase-dependent apoptosis under serum-free conditions in SH-SY5Y cells. This result is consistent with our earlier observations of mitochondrial damage and ER stress, further supporting the conclusion that cell-permeable CB1R antagonists trigger apoptotic cell death in serum-free environments.

**Figure 6.**
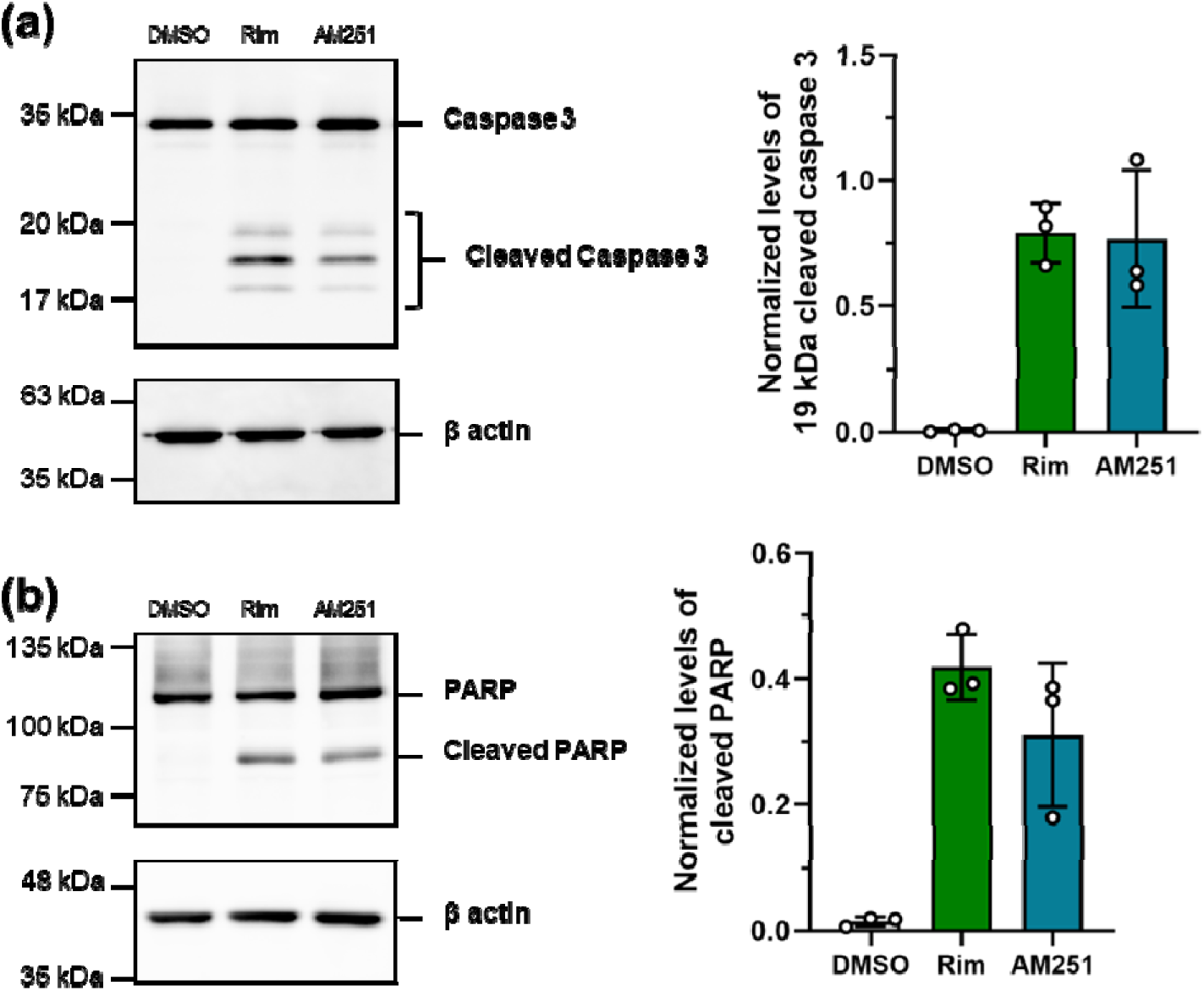
caspase-mediated apoptosis induced CB1R antagonists. SH-SY5Y Cells were treated with 10 μM rimonabant (Rim) or 10 μM AM251 for three hours. Cleaved caspase 3 (a) and cleaved PARP (b) from cell lysates were analyzed by Western blotting. The data of bar graph are shown as the mean ± SD for n = 3 (3 per independent experiment). Expression levels of 19 kDa cleaved caspase 3 and cleaved PARP were normalized by β actin expression per independent experiment.

## Discussion

In this study, we demonstrated that cell-permeable CB1R antagonists, rimonabant and AM251, induced cell death in SH-SY5Y cells under serum-free conditions, while the cell-impermeable antagonist hemopressin did not. Notably, these antagonists had no effect on cell viability in a serum-containing growth medium. At the onset of cell death under serum-free conditions, we observed mitochondrial swelling characterized by network fragmentation and cristae reduction. Moreover, we found upregulation of EIF2AK3/eIF2α/ATF4/CHOP signaling pathway, a well-known ER stress response, along with increased levels of cleaved caspase 3 and PARP. Collectively, our results suggest that neurotoxicity induced by cell-permeable CB1R antagonists under serum-free conditions is mediated by caspase-dependent apoptosis, accompanied by mitochondrial dysfunction and ER stress.

While many studies have demonstrated the neuroprotective effects of CB1R antagonists (Chan et al., 1998; Pegorini et al., 2006; Pasquariello et al., 2009; Liu et al., 2010; Tomiyama & Funada, 2014), our findings align with a subset of research that suggests potential neurotoxic effects under specific conditions. For instance, AM251 has been shown to increase caspase 3 expression in hippocampal slices, although the specific cell types affected were not identified (Caltana et al., 2015). Additionally, rimonabant has been reported to reduce cell viability in non-neuronal cells (Malfitano et al., 2010). These studies, along with our current results, highlight a complex role for CB1R antagonists, where their effects may vary depending on the cellular environment and experimental conditions.

The differential effects of CB1R antagonists in serum-containing versus serum-free conditions underscore the critical role of the cellular environment in modulating cell death signals. Fetal bovine serum is widely used as a supplement for cell growth medium, as it contains various bioactive molecules that support cell survival and growth, which can alleviate environmental stress (Zhang et al., 2016; Soutar et al., 2019; Gotten et al., 2022). The absence of cell death in serum-containing media during rimonabant and AM251 treatments can be attributed to these protective serum components. Notably, the protective effect of serum is unlikely due to direct interference with CB1R antagonist activity, as evidenced by rimonabant’s effectiveness as an orally administered drug despite circulation in the bloodstream (Henness et al., 2006). This suggests that the serum-mediated protection observed in our study is more likely due to its ability to enhance cellular resilience rather than directly neutralizing the antagonists. These observations highlight the importance of considering the cellular microenvironment when evaluating pharmacological agents’ effects on neuronal health and survival, suggesting that CB1R antagonists’ potential neurotoxicity may be more pronounced under specific conditions.

Serum deprivation is well-established to increase cellular sensitivity to apoptosis. This heightened sensitivity is primarily mediated through the integrated stress response (ISR), which activates the GCN2/ eIF2α/ATF4 (Ye et al., 2010; Pakos-Zebrucka et al., 2016). Our findings demonstrate that rimonabant or AM251 treatment activates the EIF2AK3/eIF2α/ATF4/CHOP signaling pathway, a key component of the ISR signaling pathway (Pakos-Zebrucka et al., 2016). In serum-free conditions, cells are likely primed for apoptosis due to the activated ISR, making them more susceptible to additional stressors. Thus, CB1R antagonists may trigger apoptosis more readily in these sensitized cells by further amplifying the eIF2α/ATF4/CHOP signaling cascade, effectively lowering the threshold for apoptosis induction. This synergistic effect between serum deprivation and CB1R antagonism could explain the observed cell death in our serum-free experiments.

Calcium overload is a key factor in mitochondrial dysfunction and ER stress (Bravo et al., 2011; Panda et al., 2021). Mitochondria serve as major intracellular calcium reservoirs, and excessive calcium accumulation triggers the mitochondrial permeability transition pore (mPTP) opening, leading to mitochondrial swelling and apoptosis (Lemasters et al., 2009; Orrenius et al., 2015). Additionally, calcium transport from the ER to mitochondria can exacerbate ER stress, promoting further mitochondrial dysfunction (Bravo et al., 2011). CHOP has been shown to promote the expression of ER oxidoreductin 1α (ERO1α), which activates the ER calcium release channel inositol-1,4,5-trisphosphate receptor 1 (IP3R1), causing calcium leakage from the ER into the cytoplasm (Sevier & Kaiser, 2007; Rozpedek et al., 2016). Therefore, Rim and AM251 may induce cytosolic calcium overload, triggering caspase-dependent apoptosis via mitochondrial and ER dysfunction.

Our observation that cell-permeable CB1R antagonists exhibited cytotoxicity while hemopressin did not, suggests that intracellular CB1Rs may contribute to the cytotoxicity. Recent studies have demonstrated that CB1R is expressed not only on the cell plasma membrane but also on the mitochondrial outer membrane (mtCB1R) in various cell types, including neurons and astrocytes (Bénard et al., 2012; Hebert-Chatelain et al., 2016; Serrat et al., 2021). These mtCB1Rs are known to modulate mitochondrial calcium uptake from the ER (Serrat et al., 2021). Given that rimonabant and AM251 can penetrate cells, they may antagonize mtCB1Rs, potentially disrupting calcium homeostasis at the ER-mitochondria interface. This hypothesis is supported by a previous study showing that AM251, but not hemopressin, reversed CB1R-mediated neuroprotection against ischemia (Yang et al., 2020). Thus, the antagonism of mtCB1R by cell-permeable antagonists could be a mechanism underlying the observed neurotoxicity.

However, it is important to consider that rimonabant and AM251 might also induce neurotoxicity independent of CB1R. Previous studies have reported that rimonabant can undergo intracellular bioactivation, generating toxic reactive metabolites that covalently bind to cellular proteins (Bergström et al., 2011; Foster et al., 2012). The ER has been identified as a primary target of these reactive drug metabolites (Huang et al., 2022). Therefore, it is possible that the reactive metabolites of rimonabant and AM251 bind to ER proteins, leading to ER stress-mediated apoptosis. This alternative pathway could explain the observed ER stress in our study and suggests that the neurotoxic effects of these compounds may result from a combination of CB1R-dependent and independent mechanisms. Future research using CB1R knockout models or more selective CB1R antagonists could help elucidate the relative contributions of these pathways to the observed neurotoxicity.

In conclusion, our study demonstrates that cell-permeable CB1R antagonists, rimonabant and AM251, induce caspase-dependent apoptosis through mitochondrial dysfunction and ER stress under serum-free conditions in SH-SY5Y cells. Importantly, the presence of serum mitigates these neurotoxic effects, highlighting the critical role of environmental factors in modulating CB1R antagonist-induced cellular responses. These findings suggest that CB1R antagonists may have neurotoxic potential under specific conditions, such as cellular stress or nutrient deprivation. Further research is needed to fully elucidate the mechanisms underlying this conditional neurotoxicity and to assess its relevance *in vivo*. This work contributes to our understanding of CB1R antagonist actions and may have implications for their therapeutic use in various neurological conditions.

## Acknowledgement

The authors are grateful to A. L. Sugawara and M. Oishi for English proofreading and doble-blind classification of mitochondrial cristae morphology, respectively. K.M. is supported by JSPS DC1 Research Fellowships for Young Scientists (22KJ2960). This work was supported by JSPS KAKENHI, Grant Numbers 19K20196 (K.K.), 19K24693 (C.N.), 20K06730(K.O.), 22K17815 (K.K.), and 23K06239(K.O.).

## Author contributions

K.M. carried out data curation, formal analysis, investigation and visualization. A.T. and K.O. supported K.M.’s data curation in SEM observation of mitochondria and array topography of mitochondria. K.Y. and T.U. supported K.M.’s data curation in phosphor-proteomics analyses. K.M. wrote the original draft. C.N. and K.K. designed the conceptualization and methodology. Y.K., K.O., C.N. and K.K. reviewed and edited the manuscript. K.O., T.U., C.N., T.A., and K.K acquired funding and offered resource. C.N., T.A., and K.K administered and supervised the project.

## Supplemental file captions

**Movie S1**. 3D reconstruction of mitochondrial shape in a single SH-SY5Y cell in untreated group. Colored particles, mitochondria; reddish purple, cell. Scale bars: 10 μm.

**Movie S2**. 3D reconstruction of mitochondrial shape in a single SH-SY5Y cell in rimonabant treatment group. Colored particles, mitochondria; reddish purple, cell. Scale bars: 10 μm.

