## Supplemental Figures for "Mitochondrial Dysfunction and ER Stress in CB1 Receptor Antagonist-Induced Apoptosis in Human Neuroblastoma SH-SY5Y Cells"

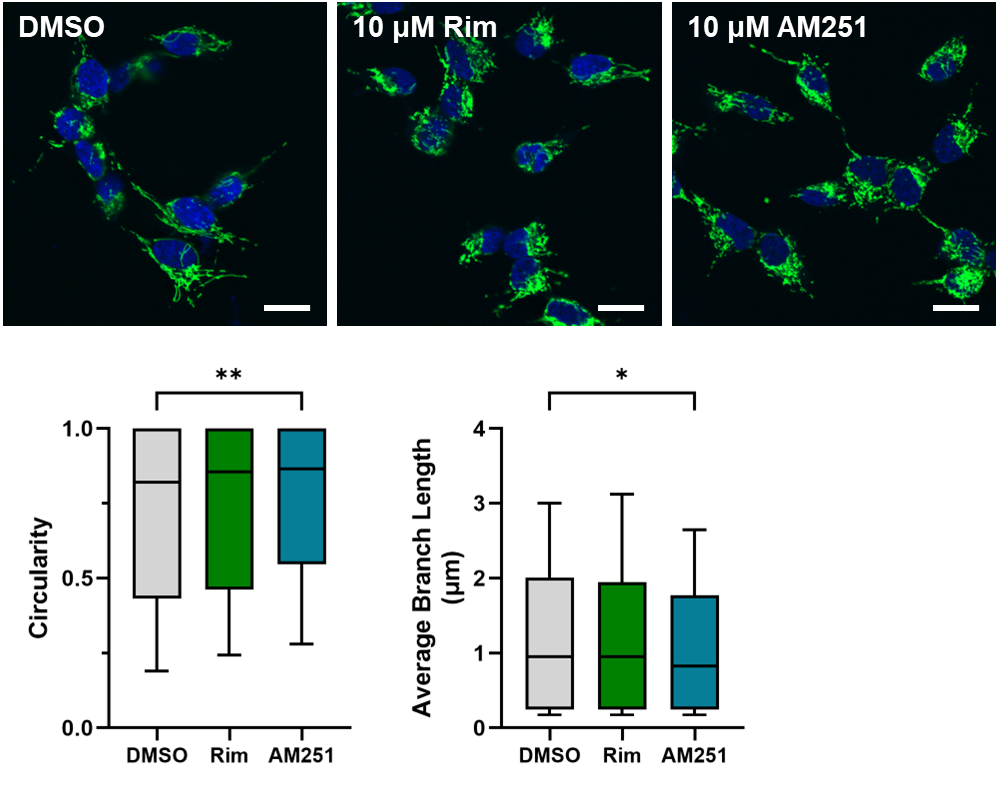

**Supplemental Figure 1.** Mitochondrial morphology of SH-SY5Y cells treated with cell-permeable CB1R antagonists under growth conditions. Cells were treated with 10 μM rimonabant (Rim) or 10 μM AM251 in a growth medium for one hour. Green, Tom20; blue, DAPI. Scale bar: 20 μm. Mitochondrial morphology was analyzed using ImageJ. Data were shown as Box and whiskers with 10–90 percentile. DMSO: 53 cells; Rim: 57 cells; AM251: 55 cells. Data were analyzed by Kruskal-Wallis ANOVA with Dunn’s multiple comparison test. * or ** indicate *P* < 0.05 or 0.01, respectively.

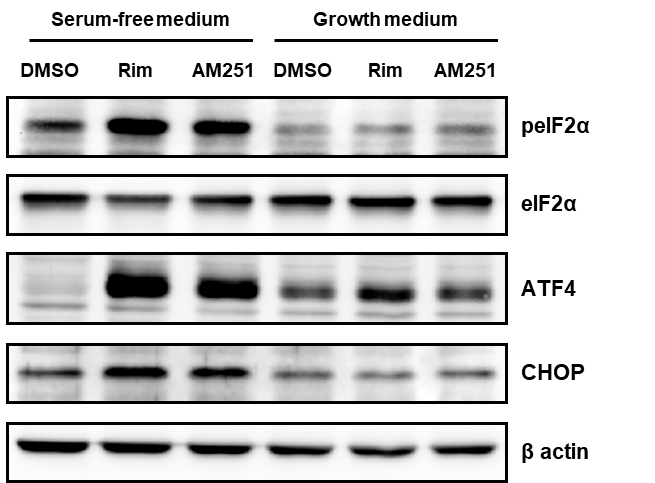

**Supplemental Figure 2**. The effect of cell-permeable CB1R antagonists on the expression of peIF2α/ATF4/CHOP under growth conditions. SH-SY5Y Cells were treated with 10 μM rimonabant (Rim) or 10 μM AM251 in a serum-free medium or a growth medium for three hours. The expression of peIF2α/ATF4/CHOP were analyzed from cell lysates by Western blotting.
